# Adult caudal fin shape is imprinted in the embryonic fin fold

**DOI:** 10.1101/2024.07.16.603744

**Authors:** Eric Surette, Joan Donahue, Stephanie Robinson, Deirdre McKenna, Crisvely Soto Martinez, Brendan Fitzgerald, Rolf O. Karlstrom, Nicolas Cumplido, Sarah K. McMenamin

## Abstract

Appendage shape is formed during development (and re-formed during regeneration) according to spatial and temporal cues that orchestrate local cellular morphogenesis. The caudal fin is the primary appendage used for propulsion in most fish species, and exhibits a range of distinct morphologies adapted for different swimming strategies, however the molecular mechanisms responsible for generating these diverse shapes remain mostly unknown. In zebrafish, caudal fins display a forked shape, with longer supportive bony rays at the periphery and shortest rays at the center. Here, we show that a premature, transient pulse of *sonic hedgehog a (shha)* overexpression during late embryonic development results in excess proliferation and growth of the central rays, causing the adult caudal fin to grow into a triangular, truncate shape. Both global and regional ectopic *shha* overexpression are sufficient to alter fin shape, and forked shape may be rescued by subsequent treatment with an antagonist of the canonical Shh pathway. The induced truncate fins show a decreased fin ray number and fail to form the hypural diastema that normally separates the dorsal and ventral fin lobes. While forked fins regenerate their original forked morphology, truncate fins regenerate truncate, suggesting that positional memory of the fin rays can be permanently altered by a transient treatment during embryogenesis. Ray finned fish have evolved a wide spectrum of caudal fin morphologies, ranging from truncate to forked, and the current work offers insights into the developmental mechanisms that may underlie this shape diversity.

## Introduction

The development and regeneration of biological shapes requires precise deployment and temporal interpretation of spatial signals (1,2); developmental shifts that alter ultimate shape can profoundly impact the function of an organ, with disordered or adaptive effects (3). The homocercal caudal fin is a major evolutionary innovation of teleosts (ray finned fish), and shows an elegant external skeletal structure that is complex enough to be developmentally informative, yet simple enough that essential aspects of form may be mechanistically disentangled. The shape of the external caudal fin is primarily derived from the difference in length between the outer dorsal and ventral (peripheral) and central fin rays. This overall shape varies considerably across species with different swimming ecologies, and the shape of the fin corresponds to different hydrodynamic tradeoffs (4–6). Across the spectrum of caudal fin diversity, there are two classes of fin shapes. Some teleosts possess triangular–truncate–shapes (e.g. in medaka, trout) with central fin rays as long as or longer than the peripheral rays; this morphology provides a relatively large surface area for efficient acceleration (5–7). Other teleost groups have evolved a forked fin shape (e.g. tuna, carp), with central rays that are relatively shorter than peripheral rays; this forked shape reduces the overall fin surface and is believed to maximize either efficient cruising or stability (4,5,8,9). The zebrafish caudal fin exhibits a distinctly forked shape and, along with the other fins, is intensively studied as a model for skeletal growth regulation and regeneration (e.g. see 10–12).

The zebrafish caudal fin is characterized by mirror-image symmetry of the rays, reflected around the central hypural diastema, a cleft that separates the central-most skeletal elements (13–15). This external symmetry contrasts with the highly asymmetric caudal fin endoskeleton, where most fin rays are supported by hypurals—modified ventral spines (16,17). During development, central caudal fin rays appear first, ossifying in pairs around the hypural diastema and ventral to the notochord, with peripheral rays appearing later in sequence (13,18). The notochord flexes upward as the fin develops, re-orienting the organ from ventral to posterior and ultimately establishing the dorsoventrally-symmetrical organ (13,15,19). Zebrafish fins are highly regenerative, and the caudal fin can regrow to its original size and forked shape within weeks of amputation (20–22).

The outgrowth of the caudal fin is initiated by pulses of cell proliferation at the distal end of the caudal fin fold mesenchyme (23,24). Skeletal precursors differentiate into osteoblasts that secrete the mineralized collagenous matrix that forms the fin rays (25,26). Signaling pathways such as Wnt, Shh, and BMP, among others, regulate the timing of skeletal differentiation, proliferation, and migration throughout the median fin fold as the organ develops and regenerates (12,27–32). Previous studies have identified several factors that act at the organ level to shape and pattern the appendage: Hox factors initially govern fin ray length, number and identity (14,33); ion channels and gap junctions govern fin ray growth by modulating tissue-level bioelectricity (34–37); thyroid hormone and osteoclast activity regulate patterning of the rays and location of ray branches (38,39). Disrupting any of these pathways profoundly disrupts the phenotype of the fin; notably however, the forked shape of the organ remains remarkably consistent even if length or skeletal patterning (or both) are disrupted (38,40). Unlike length and skeletal patterning, the developmental pathways that regulate caudal fin shape remain unresolved.

In tetrapod limbs, Shh establishes anteroposterior axis patterning, regulating limb growth and posterior skeletal identities (41–44), making the pathway a strong candidate to regulate caudal fin shape. Although *shh* is not expressed in the early caudal fin fold primordium (45), the morphogen is produced later as the skeleton develops, initially expressed along the rays and eventually localizing to the growing distal tips (45–47). During both development and regeneration, the Shh pathway promotes ray branching by trafficking pre-osteoblasts distally with migrating basal epidermis (29,47,48). However, despite the involvement of Shh in patterning vertebrate appendages and specifically regulating fin ray growth, the pathway has not previously been shown to contribute to establishing the shape of the caudal fin. Here, we demonstrate that modulating Shh in the embryonic fin fold is capable of producing a novel caudal fin shape, shifting zebrafish from a forked to a truncate caudal fin morphology.

## Results

### *Transient, premature* shha *overexpression during embryonic development alters shape and pigmentation of the caudal fin*

In wild-type (WT) zebrafish caudal fins, the shortest central rays are ∼65% the length of the longest peripheral rays, creating a forked shape (**Fig. 1A-B, E**). To investigate the role of Shh in the development of this shape, we used the *hsp70l:shha-EGFP* transgenic zebrafish line (49) to drive transient, precocious *shha* overexpression by heat shock before any of the caudal fin skeleton begins to form: 2 days post-fertilization (dpf); hereafter this treatment is referred to as “*shha* pulse”. After treatment with a *shha* pulse, the central rays grew to nearly the same length as the peripheral rays (often >85%), resulting in a truncate fin shape reminiscent of the caudal fins of medaka or killifish (**Fig. 1C-E)**. Notably, while the shape of the fin was changed, the overall size (as measured by the longest peripheral rays) was unchanged (**Fig. S1**). Fin shape showed no correlation with sex (**Fig. S2**). Fish treated with a *shha* pulse exhibited fewer principal rays, varying between 8 and 17 instead of the typical 18 (**Fig. 1F**), and showed a striking loss of the hypural diastema (**Fig. 1D, G**).

**Fig. 1:**
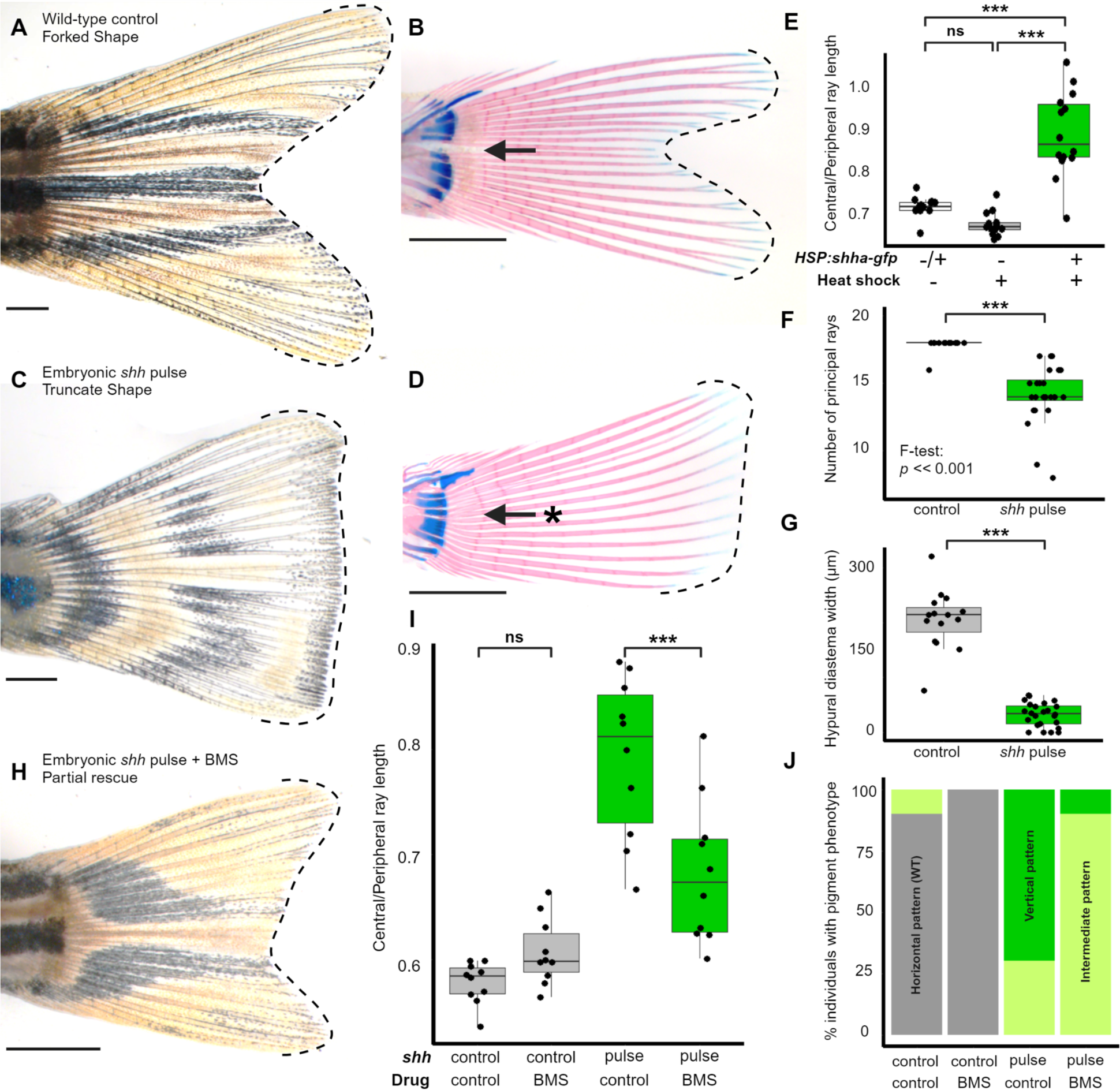
Pulse of premature *shh* during early fin fold development disrupts adult caudal fin shape. (*A-B*) Caudal fins of control zebrafish and (C-D) transgenic zebrafish subjected to transient *shh* overexpression 2 dpf (*shh* pulse). (*B and D*) are cleared and stained caudal fins from juvenile zebrafish. Dashed outlines indicate the overall shape of the fins. Arrow indicates the location of the hypural diastema separating the dorsal from ventral lobes in B; asterisk indicates the absence of the diastema in (*D*). (*E*) The inheritance of the *hsp70l:shha-eGFP* transgene and the activation of the promoter by heat shock are both necessary in order to induce the truncate fin phenotype. Significance determined by ANOVA followed by Tukey’s post hoc test. An embryonic *shh* pulse (*F*) increases the number and variance of principal fin rays and (*G*) causes a loss of the hypural diastema. Significance determined using Welch’s two-sample T-tests. (*H-J*) Treatments with the Smoothened inhibitor BMS-833923 after *shh* pulse can partially rescue both (*I*) forked fin shape and (*J*) horizontal stripes of pigmentation. Significance determined by ANOVA followed by Tukey’s post hoc test. Scale bars, 1 mm.

To determine if the phenotypic effects of a *shha* pulse were mediated through the canonical Shh signaling pathway (29,48,50), we inhibited the Shh effector Smoothened with the antagonist drug BMS-833923, 24 and 48 hours after the global *shha* pulse (48,51,52). This inhibition partially rescued the wild-type forked fin shape to *shha* pulsed fish (**Fig. 1H-I**), demonstrating that the truncate phenotype is acquired through disruption of canonical Shh signaling.

The *shha* pulse also caused a dramatic shift in pigment pattern: truncate fins developed stripes organized in vertical arches rather than in the stereotypical pattern of horizontal stripes (**Fig. 1A, C**). The pigment pattern induced by *shh* pulse is reminiscent of the vertical bars on the fins of certain species with evolved truncate fins, including clownfish (53) and some Corvis wrasses including *Corvis flavovittata*, which develop horizontal stripes on the body and vertical pigmentation on the rounded fin (D. Parichy, personal communication; (54,55)). This pigment disruption in the *shh-*pulsed group was partially rescued by treatment with BMS-833923, as well (**Fig. 1H, J**).

### *Transient embryonic* shha *overexpression alters fin shape in a dose-dependent, local manner*

To discern the developmental window during which the *shha* pulse induces truncate fin development, we heat-shocked embryos and larvae at different days post-fertilization.

Transgenic embryos heat-shocked at 2 or 3 dpf, developed truncate fins, but not at 4 dpf or later (**Fig. 2A**). We note that the effect of the heat-shock promoter diminishes at later stages of development, which may contribute to the observed critical window (**Fig. S3A**). We then quantified the duration of excess *shha* following the 2-dpf pulse using RT-qPCR. Six hours after heat shock*, shha* mRNA levels increased approximately 10-fold and returned to control baseline by 24 hours after treatment (3 dpf; **Fig. 2B**). GFP fluorescence remained detectable for 4 days after induction (6 dpf; **Fig S3B**), consistent with the stability of the c-terminal Shh-GFP fusion protein that remains in the cytoplasm following cleavage of the Shh protein and secretion of the active n-terminal fragment (49). In WT larvae, the earliest detected Shh signaling activity in the caudal fin fold coincides with fin ray emergence, typically around 8 dpf (13,45,56), two days after transgenic Shh-GFP levels have returned to baseline in the *shha*-pulsed larvae.

**Fig. 2:**
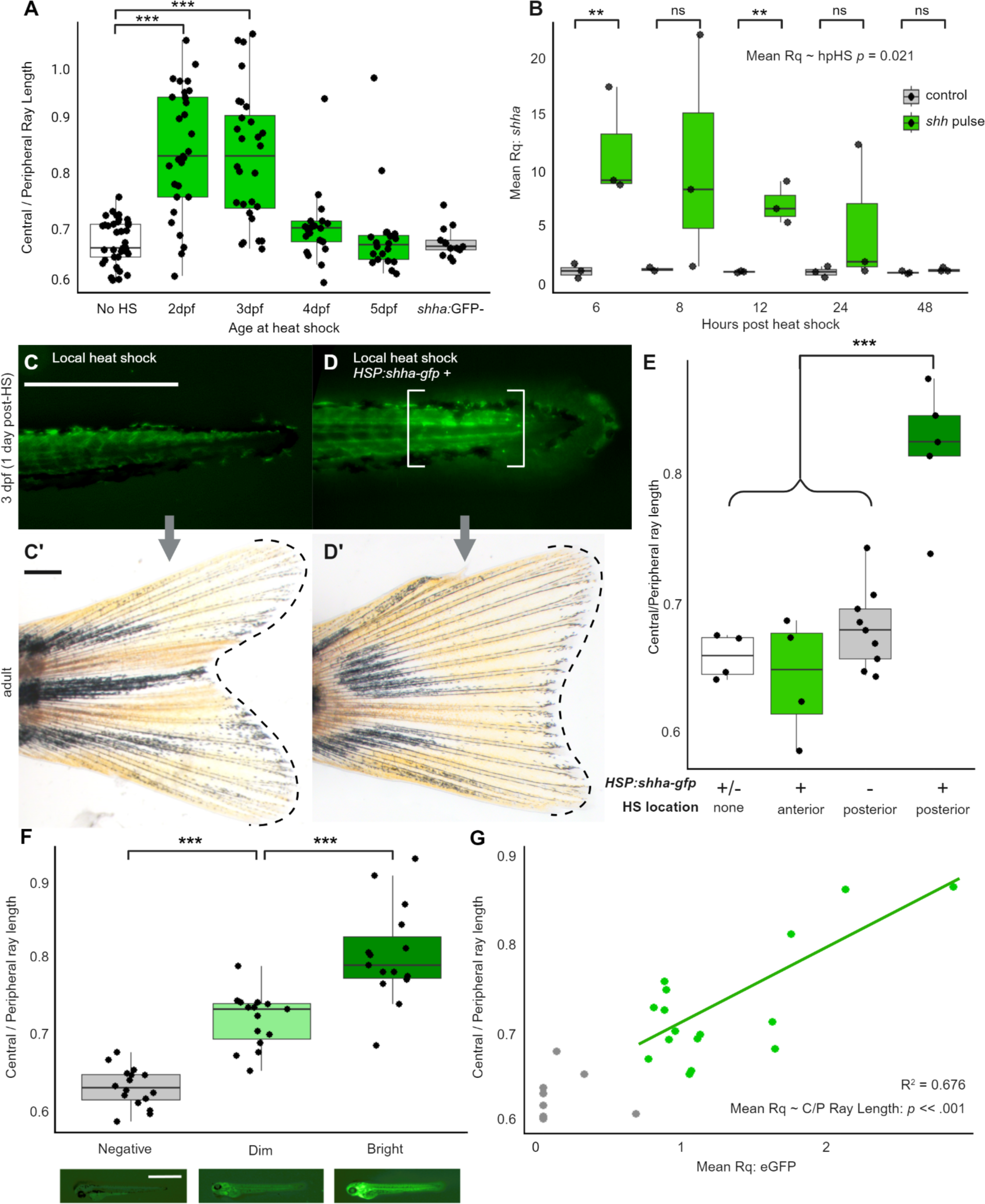
Transient embryonic *shh* pulse disrupts caudal fin development in a local and dose-dependent manner. (*A*) *shh* pulse results in truncate fin development when zebrafish are heat shock induced on 2 or 3 dpf. Significance determined by ANOVA followed by Tukey’s post hoc test. (*B*) Overexpression of *shh* during hours following heat shock. Significance determined using Welch’s two-sample T-tests, and the correlation between readout and time following heat shock determined by linear-mixed effects model. (*C-E*) Locally induced *shha* pulse is sufficient to induce truncate phenotype. (C-C’) Embryo subjected to local posterior heat shock at 2 dpf did not show GFP fluorescence and grew into an adult with a forked fin. (D-D’) Local posterior heat shock induced GFP in transgenic embryo (brackets), which grew into an adult with a truncate fin.Scale bars, 500µm. (*E*) Local posterior heat shocks in transgenic embryos are capable of inducing truncate fin shape. Inducing local *shh* pulse in the anterior of the embryo produces no change in fin shape. Significance determined by ANOVA followed by Tukey’s post hoc test. (*F*) Fish sorted by relative brightness of GFP expression 1 day after whole-body HS (4 dpf) show different caudal fin shapes as adults. Shown below the graph are representative images of individuals in each brightness category. Significance determined by ANOVA followed by Tukey’s post hoc test. Scale bar, 1 mm. (*G*) Quantified copy number of GFP transgene amplified from genomic DNA correlates with caudal fin shape. Significance between mean Rq and caudal fin shape is determined by linear-mixed effects model.

As a global *shha* pulse induced a truncate caudal fin phenotype, we predicted that *shha* overexpression solely at the posterior end of the tail would similarly produce a truncate fin. We locally activated the *hsp70l:shha-gfp* transgene using local heat shock (57) at two anteroposterior locations along the body axis. As predicted, only localized Shh activation at the posterior end of the tail (adjacent to the region where fin rays will develop (13)) was sufficient to induce truncate fin development (**Fig. 2C–E**).

We asked if the severity of the truncate phenotype correlated with the amount of *shha* transgene activation. Indeed, we found that GFP brightness following heat shock (**Fig. 2F**) as well as the quantity of genomic *gfp* (quantified by qPCR; **Fig. 2G**) each predicted the severity of the truncate phenotype. These relationships suggest that greater abundance of genomic *shha- gfp* acts in a dose-dependent manner to induce the aberrant fin shape.

### Caudal fin shape is established by regional differences in cell proliferation and growth rates

To examine the skeletal basis of the *shha* pulse-induced fin abnormalities, we tracked fin ray ossification and hypural chondrogenesis throughout larval development (**Fig. 3A–B**). In control larvae, hypurals appear from anterior to posterior, and fin rays appear sequentially in pairs around the hypural diastema from central to peripheral (see (13,15)). In *shh*-pulsed fish, the hypural complex was malformed and lacked a diastema as soon as hypurals appeared (**Fig. 3A’-B’**), while fin ray growth was delayed (**Fig. 3C**).

**Fig. 3:**
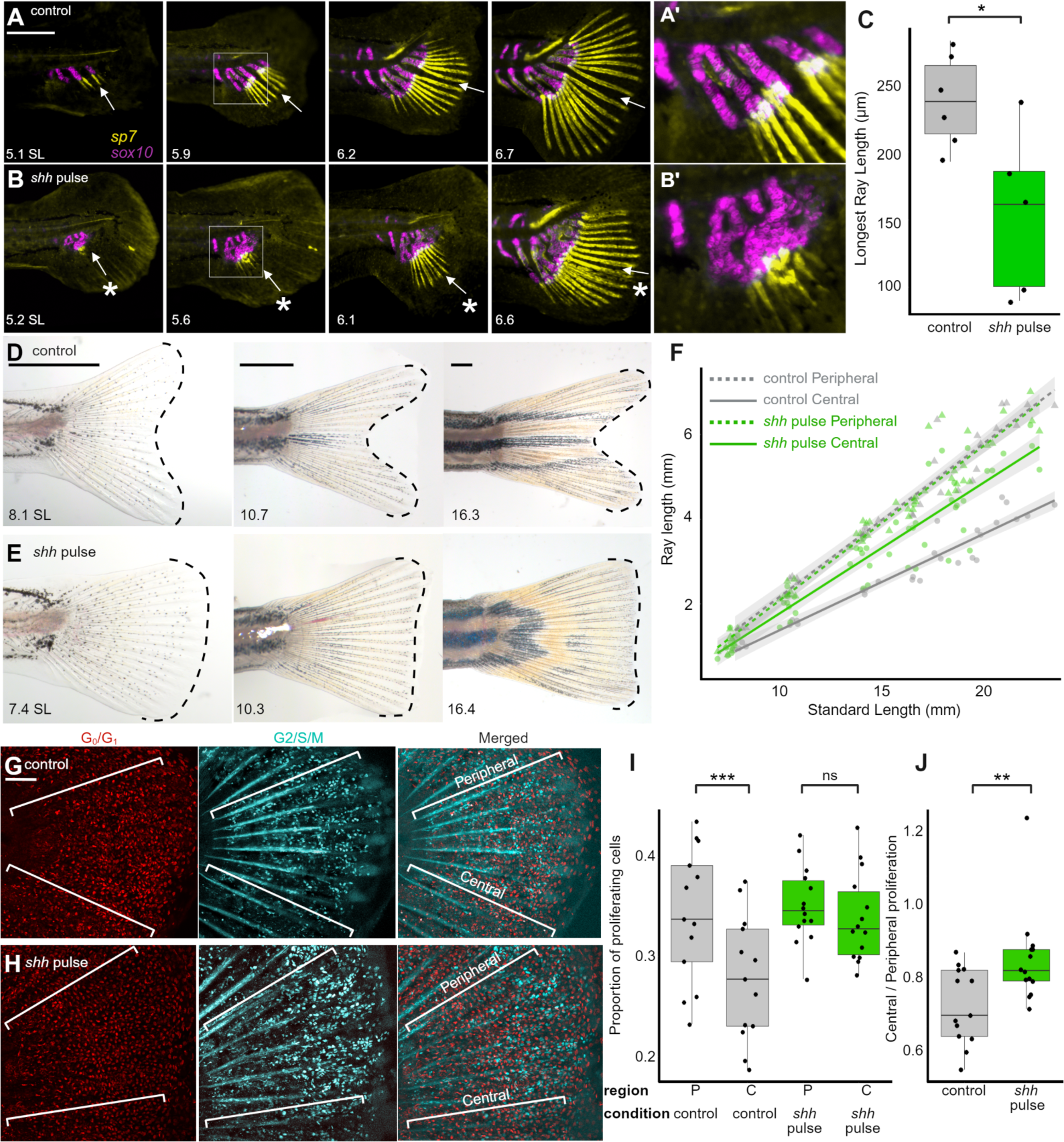
Divergent fin shape is accompanied by disruptions in skeletal growth and presaged by differences in regional cell proliferation. Development of the caudal skeleton in (A) control and (B) *shh* pulsed larvae. *sp7* reporter-expressing osteoblasts shown in yellow; *sox10* reporter-expressing chondrocytes shown in magenta. Arrow indicates the location of the hypural diastema separating the dorsal from ventral lobes in A; asterisk indicates the absence of the diastema in B in both the ossified rays and the cartilaginous hypural complex (*). Bar, 500 uM. A’ and B’ show higher magnification images of boxed areas. (*C*) In these early stages of fin development (∼6.0 SL), the length of the longest rays is less in *shh-*pulsed larvae than in control siblings. Significance determined using Welch’s two-sample T-tests. (*D*) control (*E*) and *shh* pulsed sibling caudal fin growth from 14 to 36 dpf. Dashed lines indicate the distal edge and overall shape of the fins. Scale bars, 500 µm. (*F*) The emergence of WT forked fin shape (gray lines) is the result of a lower growth rate in central rays (solid lines) relative to peripheral rays (dashed lines). Following embryonic *shh* pulse (green lines), central rays exhibit increased growth rates throughout development while peripheral rays retain a WT growth trajectory. (*G-H*) Dual *Fucci* reporter showing non-proliferating cells in red and cells in G2, S or M phase in cyan in the dorsal lobe of caudal fin folds of (*G*) control siblings and (*H*) larvae that experienced *shh* pulse. Bar, 200µm. (*I*) In control developing forked fins, proliferation is relatively lower in central regions, while *shh* pulse causes increased proliferation in central regions of the developing truncate fin Significance is determined by ANOVA followed by Tukey’s post hoc test. (*J*) Across the entire organ, proliferation becomes more uniform following *shh* pulse (closer to 1.0) compared to WT. Significance determined by Welch two-sample T-test. Difference between central / peripheral proliferation comparing WT to *shh* pulse is still significant when outlier in *shh* pulse group is removed.

We asked whether the truncate phenotype induced by *shha* pulse involved a change in the growth rate between central and peripheral rays. In control caudal fins, central rays grow slower than peripheral rays, causing the forked shape to become progressively pronounced as the fish grow (20) (**Fig. 3D**, **Fig. S1D**). In *shha*-pulsed caudal fins, peripheral rays grow at indistinguishable rates from control siblings. Strikingly however, *shha-*pulsed individuals showed 35% faster central ray growth throughout juvenile development compared to their control siblings (**Fig. 3E–F**).

We tested whether these changes in linear growth rates of rays following a *shha* pulse would correspond to altered rates of regional cell proliferation. We used the Dual *z-*Fucci cell-proliferation reporter transgenic line (55) to quantify proliferating cells in peripheral and central regions of control fins starting as early as ∼5.9mm SL, when peripheral and central regions show similar cell population sizes (see **Fig. S4A)**. Differences in regional cell proliferation (previously described in (24,59)) emerged during this developmental stage, with central regions of fins proliferating slower than peripheral regions (**Fig. 3G, I**). As the fish grows, increased peripheral cell proliferation rates lead to similarly larger cell population sizes in peripheral regions compared to center (**Fig. S4B**). The *shha* pulse led to increased proliferation specifically in the central region of fins (**Fig. 3H–J**) leading to larger central cell populations compared to control siblings (**Fig. S4A–B**). Peripheral proliferation levels did not change relative to controls (**Fig. 3I**). These results suggest that an embryonic *shha* pulse changes proliferation fates, evident at later larval stages during skeletogenesis, subsequently altering fin ray growth rates, changing the ultimate shape of the caudal fin.

These results led us to ask whether rays growing in different regions of the fin showed different levels of Shh pathway activation. Ptch2 is a receptor of Shh and also serves as a readout of canonical Shh activity (60), and expression of *ptch2* was elevated 12–24 hours after *shha* pulse treatment (**Fig. S5A**), before rays developed. Both *shha* and *ptch2* are concentrated at the distal tips of growing rays (28,44), with Ptch2 fluorescence (60) enriched in peripheral rays compared to central rays (**Fig. S5B, D**) (29). In contrast, *shha-*pulsed individuals showed comparatively elevated *ptch2* reporter activity in the central regions (**Fig. S5C-D**).

### *Embryonic* shha *overexpression alters tissue memory and acts independently from mechanisms that regulate fin length and ray pattern*

In control fins, the forked shape, length, and skeletal patterning of the caudal fin are rebuilt during regeneration (**Fig. 4A, C**). We asked if the *shha* pulse permanently altered memory of fin shape: indeed, amputated truncate fins restored their original truncate shape (**Fig. 4B-C**). The embryonic *shha* treatment induced not just a developmental change in positional information, but also a shift in positional memory consistent through adulthood.

**Fig. 4:**
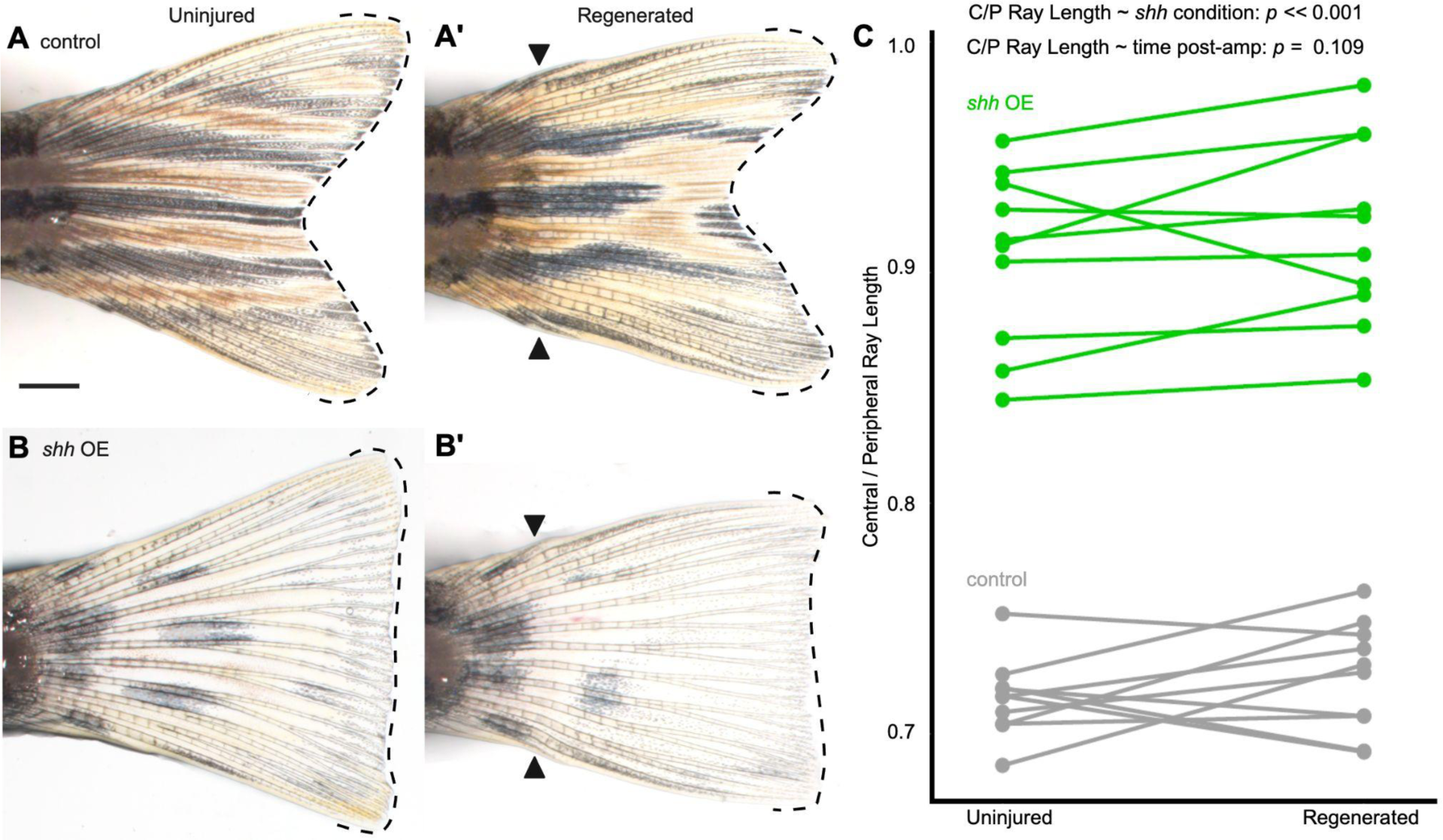
Altered memory of the adult caudal fin. (*A-B*) Control forked caudal fins (*A*) restore a forked shape 30 days after amputation. (*B*) Truncate fins restore a truncate shape after amputation. Bar, 1 mm. (*C*) Quantification showing the fin shape of each individual before and after regeneration. Significance between factors determined via a linear mixed-effects model.

Previous research characterized *longfin* and *shortfin* mutants, which show marked differences in fin size relative to SL (34,35,57); hypothyroid fish show a proximalized fin ray patterning with delayed branching (38). In these altered phenotypic contexts, the overall caudal fin shape remains forked (see **Fig. S6A, C, E**). We asked if introducing an embryonic *shha* pulse in these altered phenotypic contexts could induce a truncate phenotype. Indeed, lengthened *(lof*), shortened (*sof*), or proximalized (conditionally hypothyroid) backgrounds each produced truncate fins following the embryonic *shha* pulse (**Fig. S6B, D, F**), suggesting that length, patterning, and shape are each regulated by independent signaling pathways that may be decoupled to produce a vast range of phenotypic fin diversity.

## Discussion

We have demonstrated that the caudal fin shape of zebrafish is imprinted in the embryonic fin fold tissue, and that it may be re-patterned by excess *shh*. Shh is produced by the zone of polarizing activity (ZPA) in the posterior regions of growing limb buds, and confers posterior identity in tetrapod appendages (43). In tetrapods, loss of function to the Shh pathway prevents formation of distal limb structures (62,63), while ectopic Shh expression at the anterior border of the limb bud induces mirror-image digit duplications with posterior identities (42,43,64), with greater amounts or longer duration of Shh activation inducing supernumerary posterior digits (65,66). The posteriorizing role of Shh predates the fin-to-limb transition, a Shh-producing ZPA is both present in the pectoral and pelvic (paired) fins as well as dorsal and anal (median) fins of chondrichthyans (67,68) and bony fishes (69–71). Impairment of the Shh pathway blocks paired fin formation entirely (69,72). However, in contrast to the other median fins, no tissue with ZPA- like activity has been identified in the early caudal fin fold (44,55). Here, we have shown that an embryonic *shha* pulse in the caudal fin fold–5–6 days before *shh* is normally expressed in these tissues (45,56)–is sufficient to re-pattern shape of the organ by specifically elongating the central rays, without an obvious posteriorization effect. In contrast to phenotypes induced by excess Shh in paired appendages, the *shh* pulse in zebrafish decreases caudal fin ray number and appears to “peripheralize” the rays, with central rays now growing to lengths that match those of the peripheral rays.

Neither *shha* nor *shhb* are expressed in the posterior tail mesenchyme during zebrafish embryogenesis, and early caudal fin morphogenesis does not appear to require activation of the Shh pathway (45,56). Blocking the Shh pathway specifically during juvenile development inhibits growth of the peripheral-most non-branching rays (29), but the pathway does not appear to be required for early patterning. Indeed, although loss of fin *shh* expression blocks formation of the paired fins (above), the caudal fin (as well as the anal fin) appears to be unaffected (69).

Likewise, we found that we found that early treatment with a Smo inhibitor did not affect development of the forked caudal fin shape (see **Fig. 1I**). As previously suggested for the anal fin (70), it appears that the caudal fin utilizes regulatory signals that do not require the Shh pathway. Nonetheless, these tissues remain sensitive to premature expression of *shha*, which we show is sufficient to repattern the growing organ. Indeed, although *shh* is not expressed in the early fin fold primordium, downstream Shh effectors (including *gli3, smo and ptch2*) are expressed in these tissues (45), and these canonical effectors likely allow precocious *shha* to activate the downstream signaling cascade and reshape the fin.

In paired limbs and fins, Shh signaling functions in concert with 5’ Hox factors to establish anteroposterior patterning (73,74). In tetrapod limbs, continuous Shh expression is required for maintenance and later expression of HoxA and D cluster genes (73), and Shh inhibits the repressor form of Gli3, activating 5’ *HoxD* expression (41,75). Recent zebrafish mutant analyses demonstrate that while *hoxA* and *D* cluster mutations impair pectoral, pelvic, dorsal and anal fin development, the caudal fin remains unaffected (76). Instead, the caudal fin is regulated by *HoxB* and *C* cluster genes, specifically *hoxb13a* and *hoxc13a*, which are expressed in a region-specific manner at the posterior tail at 2-4 dpf, before the emergence of any skeletal elements (14). While double mutants for both *hoxb13a* and *hoxc13a* fail entirely to form a caudal fin, single knock out of either *hoxb13a* or *hoxc13a* reduces fin ray numbers, abolishes the hypural diastema, and alters fin shape by shortening the peripheral rays (14). The notable phenotypic similarities between single *hoxb13a* or *hoxc13a* loss-of-function phenotypes and the truncate fin phenotype presented here suggests that the *shha* pulse may act by disrupting *hox13* gene expression or regulation to repattern fin shape during development.

Desvignes et al. (13) proposed that a central organizing center establishes the hypural diastema and defines the axis of symmetry for the growing fin. According to this model, the organizing center splits the ventroposterior mesenchyme in the central endoskeleton into two plates of connective tissue, forming the diastema between the central hypurals (hypurals 2 and 3). This is followed by progressive, paired emergence of fin rays from central to peripheral; the diastema is theorized to inhibit the growth potential of the earliest developing rays at the center of the organ (13). The early pulse of *shha* may disrupt formation or activity of the diastema organizing center; in the absence of a central organizing signal, elongating rays may default to peripheral identity and growth. We found that *shha* pulse applied at 4 dpf or later was ineffective at producing a truncate fin (see **Fig. 2A**) even though these later heat shocks were capable of inducing transgene expression (see **Fig. S3B**). This suggests that the positional information that establishes relative ray length and caudal fin shape is imprinted before 4 dpf, well before *shh* is normally expressed in these tissues (45,56). Alternatively, we note it is possible that decreasing efficiency of the heat shock promoter with age may not produce sufficient *shha* to induce the positional shift at later stages.

Our data suggest that *shha* is capable—directly or indirectly—of inducing peripheral characteristics in centrally located rays, in terms of their skeletal growth and increased underlying proliferation (see **Fig 3**). In developing forked fins, peripheral regions proliferate at a higher rate than central regions (24), likely supporting the accelerated growth of these rays (see **Fig. 3G, I**). The embryonic *shha* pulse induced increased proliferation rates in the now rapidly-growing central region of the emerging truncate fin (see **Fig. 3H-J**). The Shh pathway can directly regulate proliferation rates: in parallel to its role patterning posterior limbs, Shh signaling promotes proliferation by regulating cell-cycle G1–S progression in the distal limb mesenchyme (77–79). The mesenchymal cells proliferate at higher rates in the outer regions of the distal limb bud relative to the center as the organ grows during embryonic development (80). Reduction of Shh signaling after early patterning decreases limb mesenchymal cell proliferation, leading to time-dependent progressive digit loss (78). Shh signaling affects growth by upregulating cyclin/kinase pair Ccnd1 and Cdk6 transcription through the inhibition of the repressor form of Gli3 (79). This mechanism is evolutionary conserved, as *gli3* mutants in medaka increase *ccnd1* and *cdk6* transcription, which have conserved Gli3-regulated promoters across vertebrate genomes (70). Our data presented here demonstrates that relative modulation of regional proliferation rates is associated with different adult fin shapes, providing evidence that intrinsic positional characteristics inform local growth rates across the growing fin to progressively sculpt the shape of the organ.

A pulse of *shha* in the embryonic fin can permanently alter the memory of fin shape, since the adult truncate fin phenotype was restored following amputation without additional exogenous Shh (**Fig. 4**). While modulating bioelectricity or thyroid hormone availability can shift the morphology of a regenerating fin, neither treatment is capable of altering tissue memory: fins revert to a WT morphology if the treatment (calcineurin inhibition or thyroid inhibition) is removed or rescued (30,38). Memory of fin size can be altered by inhibiting overall proliferation: temporarily inactivating an accessory subunit of the DNA polymerase alpha (*pola2*) during regeneration can permanently alter the memory of fin size (40). Notably, the forked shape is not altered by this treatment (40). Our work therefore identifies an early developmental window of caudal fin development during which positional information is imprinted that will inform both the development and the memory of organ shape.

The evolution of the externally-symmetrical homocercal caudal fin in teleosts allowed for the external skeleton to take on distinct dorsoventral functionalization (19,81–83). This morphological functionalization may have supported the diversification of caudal fin shapes across teleosts. The wide spectrum of fin shape diversity can be categorized into truncate shapes with a flat or rounded edge and forked shapes with a concave edge; truncate and forked fins each offer biomechanical advantages and tradeoffs as propulsive and stabilizing organs (5,6,19,84). Here, we have identified an experimental manipulation capable of changing the zebrafish caudal fin from a forked to a truncate shape (see **Fig. 1, 3F, S1C**). In addition to the noticeably altered external shape, induced zebrafish truncate fins consistently lacked a hypural diastema (see **Fig. 1D, G**). The hypural diastema is considered a teleostean novelty (15) (although it was independently acquired in gars (19)). The diastema has been convergently lost at least once in nearly every lineage of crown teleosts, including cusk, swamp and true eels and in derived groups of bony tongues, catfishes, cods, flatfishes and killifishes (15,19,85–87).

Notably, many of the clades that have lost the hypural diastema also show a truncate or rounded caudal fin shape, however the evolutionary relationship between hypural diastema and the shape of the fin has not been explored in detail. Our work identifies early activation of the Shh pathway or its downstream targets as capable of reshaping the external fin and abolishing the diastema, without altering length or proximodistal patterning of the organ (see **Fig S6**), suggesting developmental mechanisms that may underlie natural teleost fin diversity.

## Acknowledgements

For fish care and assistance, we thank the BC Animal Care Facility and members of the McMenamin Lab past and present. For assistance with experiments, we thank Connor Murphy and Minqi Shen, Shahid Ali and the Karlstrom Lab. For sharing fish lines, we thank Drs. Fisher, Harris, Karlstrom, Kimelman, North, Parichy, and Stankunas and their labs. For training and use of the Zeiss AxioImager Z2, we thank Bret Judson of the Boston College Imaging Facility. For statistical assistance, we thank Melissa McTernan. Funding provided by R35GM146467, a Smith Family Foundation Odyssey Award, and NSF CAREER 1845513.

## Methods

### Resource Availability

*Lead Contact*: Further information and requests for resources or reagents, including fish lines, should be directed to and will be fulfilled by the lead contact, Dr. Sarah McMenamin.

### Materials Availability

This study did not generate any new reagents or animal strains

### Data and Code Availability

● All data reported in this study will be shared by the lead contact upon request
● This paper does not report original code
● Any additional information required to reanalyze the data reported in this paper is available from the lead contact upon request.

### Experimental Model and Subject Details

#### Experimental Animals

Zebrafish were reared under standard conditions at 28°C with a 14:10 light:dark cycle. Fish were fed marine rotifers, *Artemia*, Adult Zebrafish Diet (Zeigler, Gardners PA, USA) and Gemma Micro (Skretting, Stavanger, NOR). Individuals that experienced a *shha* pulse during embryogenesis (below) experienced a slight growth delay during larval development, although caught up in size by early juvenile stages (see **Fig. S1**). Because of the early growth delay, we took care to size-match treated and control individuals and standard length (SL) are reported throughout. Note that prior to development of the hypural complex, notochord length was measured, and is referred to as SL per (18). For developmental serial imaging, siblings positive and negative for the *hsp70l:shha-eGFP* transgene were reared in individual containers so individuals could be identified. Fish line to induce *sonic hedgehog a* overexpression was *Tg*(*hsp70l:shha-EGFP*) (49). Other lines used were *Tg(sp7:GFP)b1212* (88) to visualize osteoblasts, *Tg(p7.2sox10:mRFP)* (89) for chondrocytes, *TgBAC(ptch2:Kaede)* (60), and Dual *z-Fucci* (58,90). Mutants used were *longfin^dt2^/kcnh2a* (35,91), and *shortfin^dj7e2^/cnx43* (34).

### Method Details

#### Imaging

Zebrafish were anesthetized with tricaine (MS-222, ∼0.02% w/v in system water). Anesthetized or cleared and stained (92) individuals were imaged on an Olympus SZX16 stereoscope using an Olympus DP74 camera, an Olympus IX83 inverted microscope using a Hamamatsu ORCA Flash 4.0 camera, a Leica Thunder Imager Model Organism using a sCMOS monochrome camera, or a Zeiss AxioImager Z2 using a Hamamatsu Flash4.0 V3 sCMOS camera. Identical exposure times and settings were used to compare experimental treatments and capture repeated images of fins. Images were correspondingly adjusted for contrast, brightness and color balance using FIJI (93), and compiled using BioRender.

*Sonic hedgehog overexpression*: *Tg*(*hsp70l:shha-EGFP*) crosses at 48-54 hours post fertilization (hpf) were treated with 37° heat shock for 15 minutes. 16-18 hours after treatment, individuals were screened for GFP expression as in (49). Sibling larvae that were treated with heat shock but were negative for GFP were kept as negative WT controls.

#### Localized shha overexpression

Localized induction of the heat shock promoter was performed as previously described (57). Local heat shocks were induced at 48-54hpf for 15 minutes; local GFP expression was confirmed ∼16-18 hours after the treatment.

#### Amputations

Adult caudal fin regeneration experiments were performed on adult zebrafish 27- 33 mm SL. Caudal fins were amputated from anesthetized fish under a stereoscope at the 5th ray segment using a razor blade and given 30 days to regenerate.

#### Drug Treatments

To rescue the *shha* overexpression phenotype by Shh pathway inhibition, larvae were treated either with the Smoothened inhibitor BMS-833923 (10mM stock in 100% DMSO, 0.5 µM working solution in fish water) or the vehicle control DMSO (0.5 µM in fish water). A clutch of Tg(*hsp70l:shha-eGFP*) was treated with heat shock and sorted as above, and transgenic (GFP+) and non-transgenic (GFP-) siblings were treated with either BMS or the vehicle control for 4 hours starting 16-18 hours after the heat shock. The treatment with BMS or vehicle was repeated a second time 24 hours after the first treatment. After washout, fish were reared to adulthood under standard conditions.

To induce hypothyroidism in regenerating fish, fins were amputated as above, and allowed to regenerate in 1.0mM MPI cocktail (1.0mM MMI + 0.1mM KClO4 + 0.01mM iopanoic acid, diluted in fish water) (38,53), for 21 days with drug changes every 1-2 days.

#### RT-qPCR

Larvae were reared and heat shocked as stated above. At 3dpf, larvae were sorted into positive and negative cohorts based on fluorescence, placed into Thermo Fisher’s RNAlater™ Stabilization Solution (Cat. #: AM7021), and subsequent collection from both groups proceeded until 4dpf. RNA was extracted from these samples using Zymo Research Quick- RNA™ Microprep Kit (Cat. #: R1050) and cDNA libraries synthetized using Thermo Fisher SuperScript™ IV Reverse Transcriptase (Cat. #: 18090010). Thermo Fisher PowerUp™ SYBR™ Green Master Mix was used for qPCR (Cat. #: A25741); three technical replicates and three biological replicates were run on Thermo Fisher QuantStudio™ 3 Real-Time PCR System (Cat. #: A28567). Results were analyzed using ThermoFisher Connect Platform. The primer sets used can be found in **Table S1**.

#### Proliferation quantification

Proliferation was measured in different regions of the growing fins using the Dual *z-Fucci* transgenic line (58). Four rays were measured from each fin: the second dorsal and second ventral rays (the peripheral rays) and the two center-most rays of each lobe (the central rays). The proliferation rate was measured along a line drawn through each ray and was calculated as the number of cyan-expressing cells divided by the total number of fluorescent cells (cyan plus red). Regional proliferation was calculated as the average proliferation rate of the two peripheral rays and the average proliferation of the two central rays.

#### Statistical analysis

Analyses were performed in RStudio. Data were analyzed with either Welch two-sample t-test, ANOVA followed by Tukey’s honest significant differences (using 95% family- wise confidence level), Fligner-Killeen test, or a linear mixed-effects model. In graphs showing pairwise comparisons, significance is indicated as follows: * *p* < 0.05, ** *p* < 0.01, *** *p* < 0.001.

## Supp. Figures

**Fig. S1:**
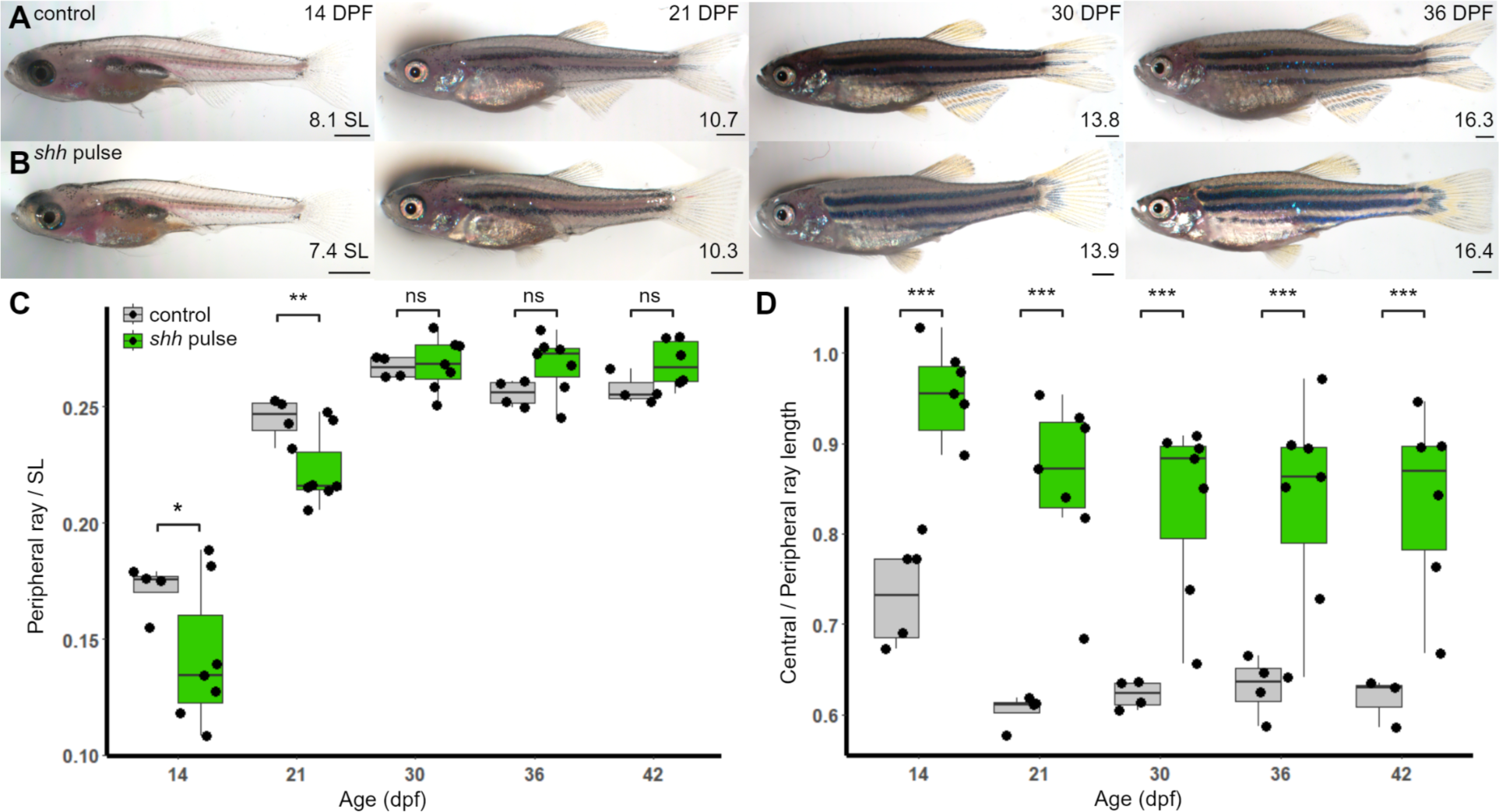
Growth of body and fins under different *shha* profiles. *A-B*) Whole body images of (*A*) sibling control and (*B*) *shh* pulse-treated fish from 14-36dpf. Scale bars, 500 µm. (*C*) The overall length of the caudal fin gfas measured by the length of the peripheral ray) relative to the standard length of the fish. By 30 dpf, truncate fins were the same size as forked fins of control siblings. (*D*) The difference in caudal fin shape between conditions is evident by 14dpf. Significance within each time point determined by Welch’s two-sample T-tests.

**Figure S2:**
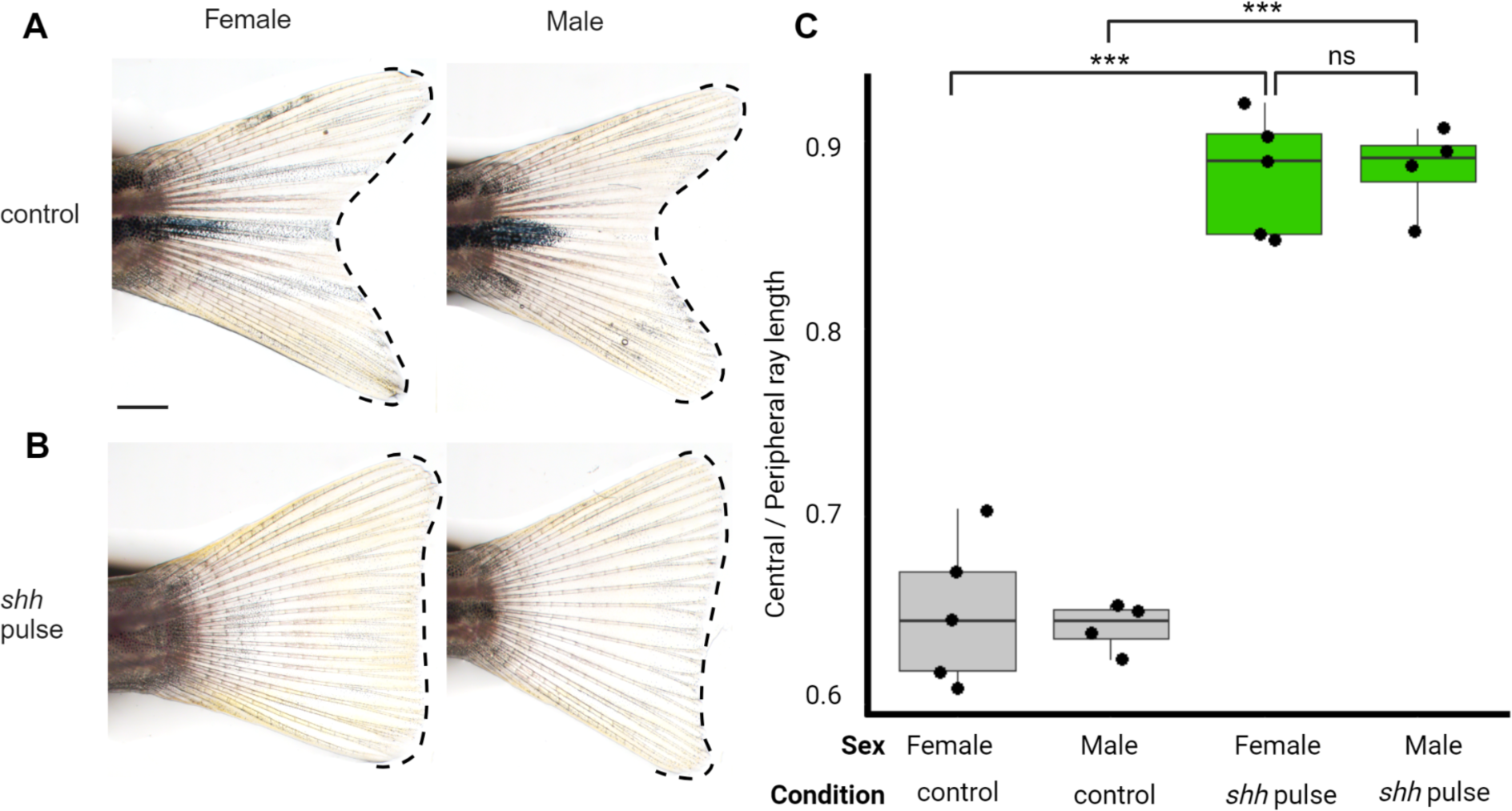
Fin shape shows no interaction with sex. Representative caudal fins of male and female (*A*) control and (*B*) *shh* pulse-treated sibling fish. Bar, 1mm. (*C*) There was no difference in fin shape between sexes in either control or *shh* pulse-treated fish. Significance determined by ANOVA followed by Tukey’s post hoc test.

**Fig. S3:**
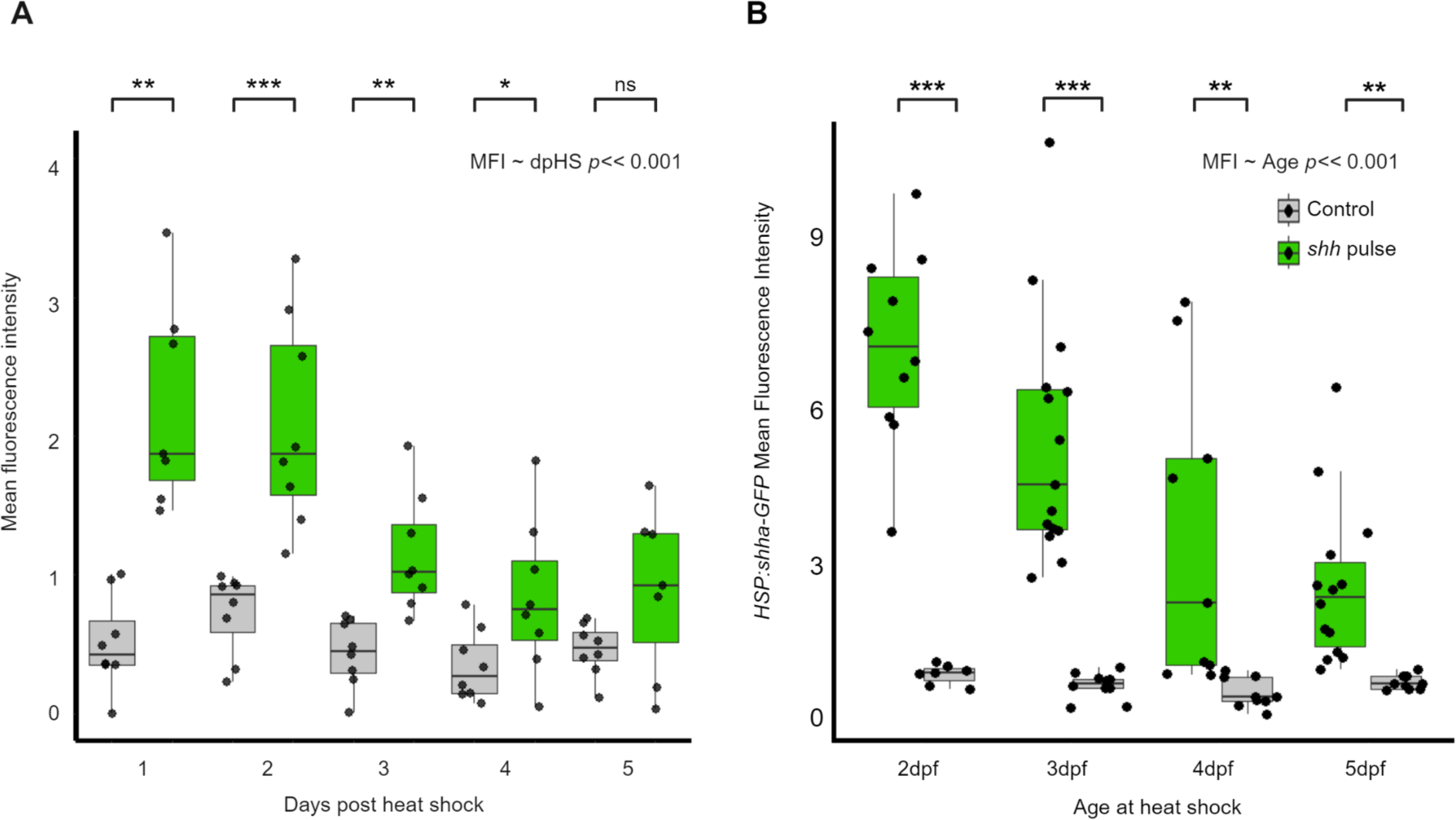
Effect of development on heat-shock promoter efsiciency and GFP perdurance following *shh* pulse (*A*) GFP fluorescence is detectable for several days following heat shock. (*B*) Amount of GFP induced decreases with later heat shocks. GFP measured by fluorescence intensity 24 h after heat shock. Significance between conditions per day determined using Welch’s two-sample T-tests, and the correlation between readout and time following heat shock / age of heat shock determined by linear-mixed effects model.

**Fig. S4:**
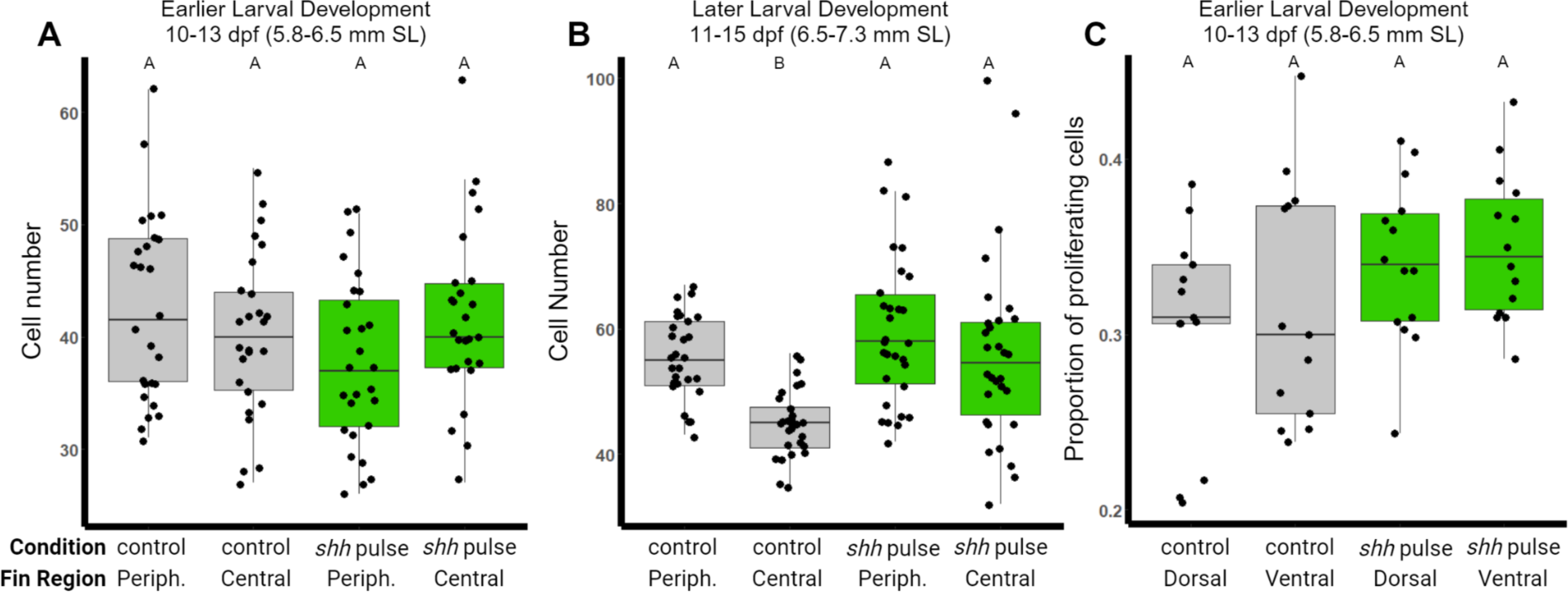
***shh* pulse leads to larger cell populations in central regions of fins.** (*A*) At earlier stages of larval development (SL = 5.8-6.5 mm) there is no difference in cell number between central and peripheral fin regions in either condition . (*B*) At later stages of larval development (SL = 6.5-7.2 mm), there is a central / peripheral differential in cell number in control individuals, but proliferation is the same in both regions of fish treated with *shh* pulse . (*C*) There is no difference in proliferation between dorsal compared to ventral fin regions . Significance determined by ANOVA followed by Tukey’s post hoc test. Statistically indistinguishable groups are shown with the same letter (threshold for significance *p* < 0.05).

**Fig. S5:**
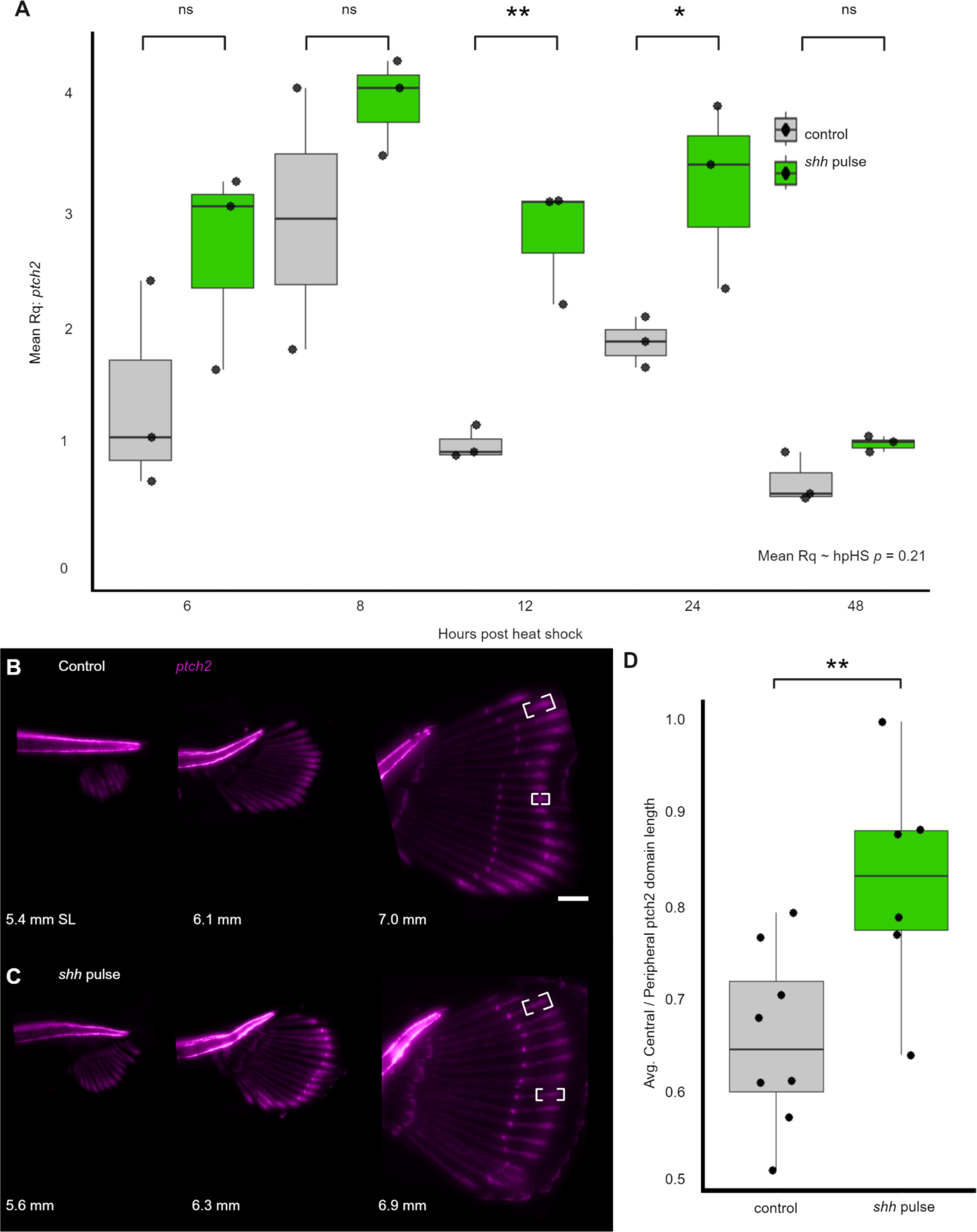
Length of *ptch2* domain correlates to relative ray length. (*A*) *shh* pulse induces moderate upregulation of *ptch2* for 24 hours following heat-shock before returning to WT levels of expression. Significance within time points determined using Welch’s two-sample T-test. Relationship between mean Rq and time following heat shock also captured by linear-mixed effects model. (*B-C*) Fluorescent image series of individual *ptch2:kaede* transgenic larvae during early caudal fin development. (*B*) In control caudal fin *ptch2:kaede* is expressed in relatively longer domains of activity in peripheral rays compared to central rays. (*C*) Following *shh* pulse, the activity domains are of similar lengths. Bar, 100µm. (*D*) The average domain length of the peripheral to the central region between conditions. Significance determined using Welch’s two-sample T-test.

**Fig. S6:**
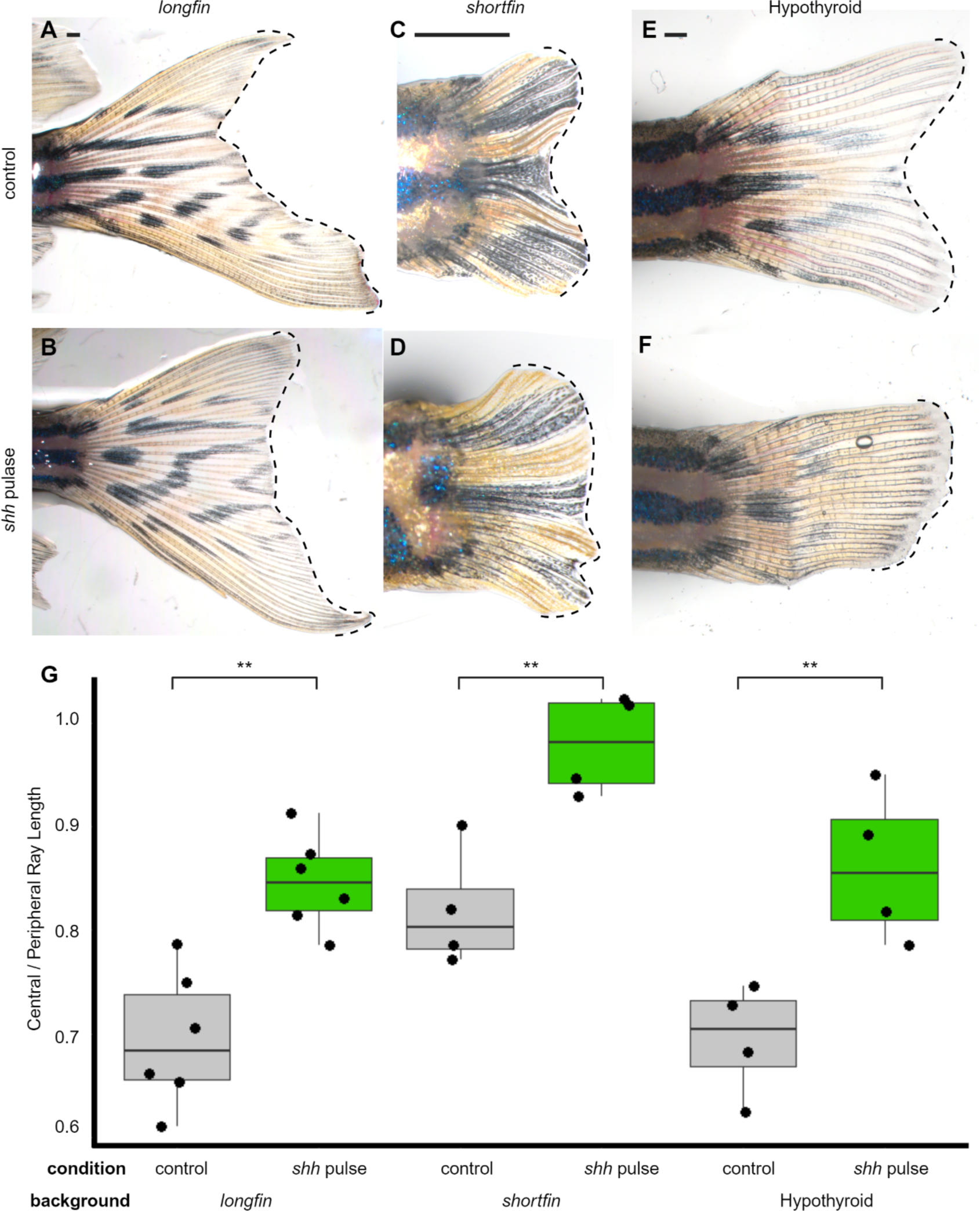
***shha* pulse induces a truncate phenotype in *longfin* and *shortfin* mutants and in hypothyroid backgrounds.** (*A, C, E*) *longfin* mutants, *shortfin* mutants, and hypothyroid zebrafish all show forked fin shape. (*B, D, F*) Treated with a *shh* pulse, truncate fin shape can be induced in all three backgrounds. (G) Quantification of fin shape in different backgrounds. Significance within each background is determined by Welch’s two-sample T-tests. Scale bars, 500µm.

**Table S1:**
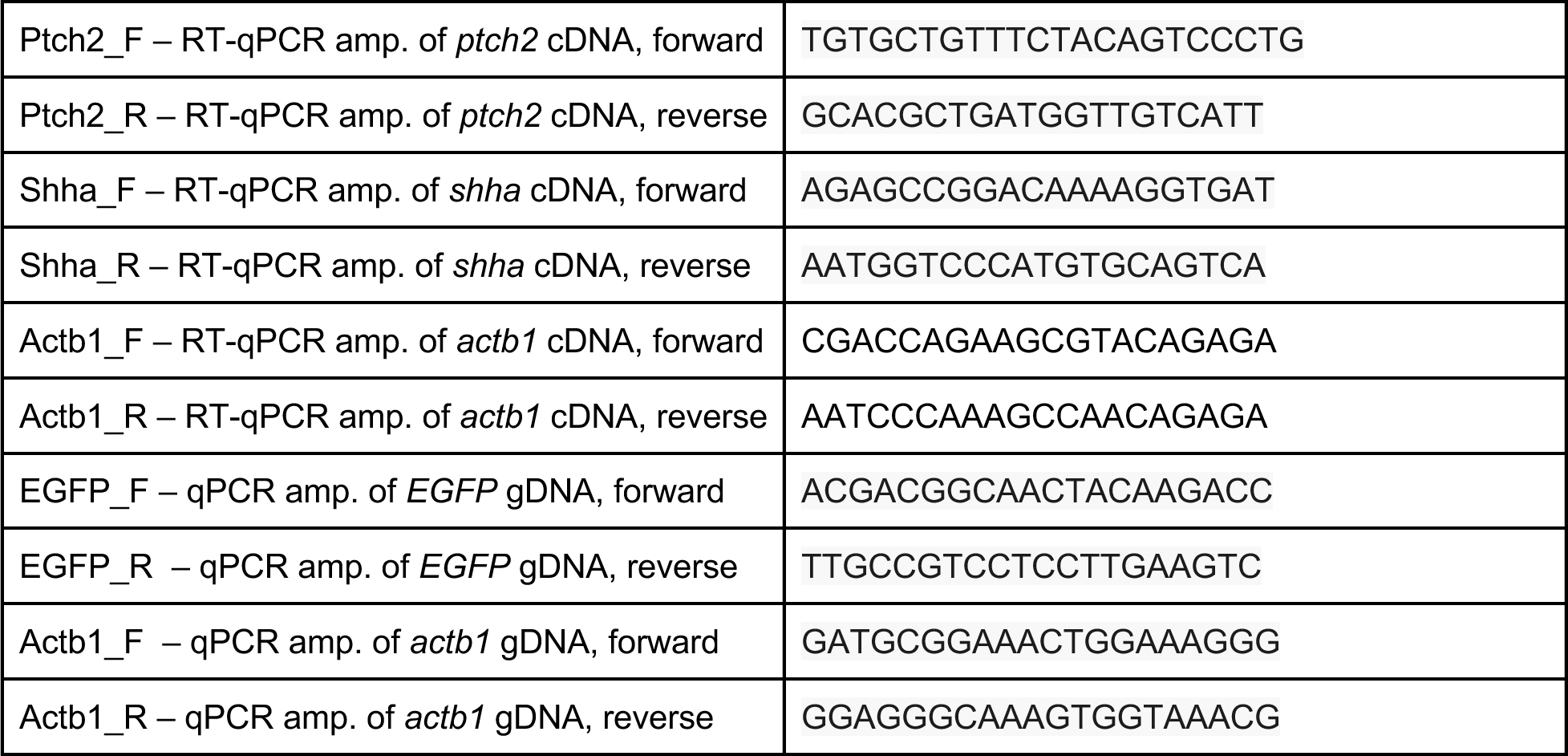
Primer sets used for qPCR and RT-qPCR.

## Reference

1. Sears K, Maier JA, Sadier A, Sorensen D, Urban DJ. Timing the developmental origins of mammalian limb diversity. genesis. 2018;56(1):e23079.

2. Wolpert L. Positional information and pattern formation in development. Dev Genet. 1994;15(6):485–90.

3. Polly PD. Chapter 15. Limbs in Mammalian Evolution. In: Chapter 15 Limbs in Mammalian Evolution. University of Chicago Press; 2008. p. 245–68.

4. Webb PW. Body Form, Locomotion and Foraging in Aquatic Vertebrates. Am Zool. 1984 Feb;24(1):107–20.

5. Tack NB, Gemmell BJ. A tale of two fish tails: does a forked tail really perform better than a truncate tail when cruising? J Exp Biol. 2022 Nov 24;225(22):jeb244967.

6. Flammang BE, Lauder GV. Caudal fin shape modulation and control during acceleration, braking and backing maneuvers in bluegill sunfish, Lepomis macrochirus. J Exp Biol. 2009 Jan 15;212(2):277–86.

7. Giammona FF. Form and Function of the Caudal Fin Throughout the Phylogeny of Fishes. Integr Comp Biol. 2021 Aug 1;61(2):550–72.

8. Flammang BE. The fish tail as a derivation from axial musculoskeletal anatomy: an integrative analysis of functional morphology. Zoology. 2014 Feb 1;117(1):86–92.

9. Liao JC. Fish swimming efficiency. Curr Biol. 2022 Jun 20;32(12):R666–71.

10. Pfefferli C, Jaźwińska A. The art of fin regeneration in zebrafish. Regeneration. 2015 May 19;2(2):72–83.

11. Henke K, Farmer DT, Niu X, Kraus JM, Galloway JL, Youngstrom DW. Genetically engineered zebrafish as models of skeletal development and regeneration. Bone. 2023 Feb 1;167:116611.

12. Harris MP, Daane JM, Lanni J. Through veiled mirrors: Fish fins giving insight into size regulation. WIREs Dev Biol. 2021;10(4):e381.

13. Desvignes T, Robbins AE, Carey AZ, Bailon-Zambrano R, Nichols JT, Postlethwait JH, et al. Coordinated patterning of zebrafish caudal fin symmetry by a central and two peripheral organizers. Dev Dyn. 2022;251(8):1306–21.

14. Cumplido N, Arratia G, Desvignes T, Muñoz-Sánchez S, Postlethwait JH, Allende ML. Hox genes control homocercal caudal fin development and evolution. Sci Adv. 2024 Jan 19;10(3):eadj5991.

15. Schultze HP, Arratia G. The caudal skeleton of basal teleosts, its conventions, and some of its major evolutionary novelties in a temporal dimension. In: Mesozoic Fishes 5-Global Diversity and Evolution. 2013. p. 187–246.

16. Cumplido N, Allende ML, Arratia G. From Devo to Evo: patterning, fusion and evolution of the zebrafish terminal vertebra. Front Zool. 2020 Jun 1;17(1):18.

17. Sanger TJ, McCune AR. Comparative osteology of the Danio (Cyprinidae: Ostariophysi) axial skeleton with comments on Danio relationships based on molecules and morphology. Zool J Linn Soc. 2002 Aug 1;135(4):529–46.

18. Parichy DM, Elizondo MR, Mills MG, Gordon TN, Engeszer RE. Normal Table of Post-Embryonic Zebrafish Development: Staging by Externally Visible Anatomy of the Living Fish. Dev Dyn Off Publ Am Assoc Anat. 2009 Dec;238(12):2975–3015.

19. Desvignes T, Carey A, Postlethwait JH. Evolution of caudal fin ray development and caudal fin hypural diastema complex in spotted gar, teleosts, and other neopterygian fishes. Dev Dyn Off Publ Am Assoc Anat. 2018 Jun;247(6):832–53.

20. Uemoto T, Abe G, Tamura K. Regrowth of zebrafish caudal fin regeneration is determined by the amputated length. Sci Rep. 2020 Jan 20;10(1):649.

21. Azevedo AS, Grotek B, Jacinto A, Weidinger G, Saúde L. The Regenerative Capacity of the Zebrafish Caudal Fin Is Not Affected by Repeated Amputations. PLOS ONE. 2011 Jul 28;6(7):e22820.

22. Sehring IM, Weidinger G. Recent advancements in understanding fin regeneration in zebrafish. WIREs Dev Biol. 2020;9(1):e367.

23. Goldsmith MI, Fisher S, Waterman R, Johnson SL. Saltatory control of isometric growth in the zebrafish caudal fin is disrupted in *long fin* and *rapunzel* mutants. Dev Biol. 2003 Jul 15;259(2):303–17.

24. Jain I, Stroka C, Yan J, Huang WM, Iovine MK. Bone growth in zebrafish fins occurs via multiple pulses of cell proliferation. Dev Dyn. 2007;236(9):2668–74.

25. Haas HJ. Studies on mechanisms of joint and bone formation in the skeleton rays of fish fins. Dev Biol. 1962 Aug 1;5(1):1–34.

26. Becerra J, Montes GS, Bexiga SRR, Junqueira LCU. Structure of the tail fin in teleosts. Cell Tissue Res. 1983 Mar 1;230(1):127–37.

27. Rabinowitz JS, Robitaille AM, Wang Y, Ray CA, Thummel R, Gu H, et al. Transcriptomic, proteomic, and metabolomic landscape of positional memory in the caudal fin of zebrafish. Proc Natl Acad Sci. 2017 Jan 31;114(5):E717–26.

28. Thummel R, Ju M, Sarras MP, Godwin AR. Both Hoxc13 orthologs are functionally important for zebrafish tail fin regeneration. Dev Genes Evol. 2007 Jun 1;217(6):413–20.

29. Braunstein JA, Robbins AE, Stewart S, Stankunas K. Basal epidermis collective migration and local Sonic hedgehog signaling promote skeletal branching morphogenesis in zebrafish fins. Dev Biol. 2021 Sep 1;477:177–90.

30. Daane JM, Lanni J, Rothenberg I, Seebohm G, Higdon CW, Johnson SL, et al. Bioelectric-calcineurin signaling module regulates allometric growth and size of the zebrafish fin. Sci Rep. 2018 Jul 10;8:10391.

31. Wehner D, Weidinger G. Signaling networks organizing regenerative growth of the zebrafish fin. Trends Genet. 2015 Jun 1;31(6):336–43.

32. Benard EL, Küçükaylak I, Hatzold J, Berendes KUW, Carney TJ, Beleggia F, et al. wnt10a is required for zebrafish median fin fold maintenance and adult unpaired fin metamorphosis. Dev Dyn.

33. Sakaguchi S, Nakatani Y, Takamatsu N, Hori H, Kawakami A, Inohaya K, et al. Medaka unextended-fin mutants suggest a role for Hoxb8a in cell migration and osteoblast differentiation during appendage formation. Dev Biol. 2006 May 15;293(2):426–38.

34. Perathoner S, Daane JM, Henrion U, Seebohm G, Higdon CW, Johnson SL, et al. Bioelectric Signaling Regulates Size in Zebrafish Fins. PLOS Genet. 2014 Jan 16;10(1):e1004080.

35. Stewart S, Le Bleu HK, Yette GA, Henner AL, Robbins AE, Braunstein JA, et al. longfin causes cis-ectopic expression of the kcnh2a ether-a-go-go K+ channel to autonomously prolong fin outgrowth. Dev Camb Engl. 2021 Jun 1;148(11):dev199384.

36. Iovine MK, Higgins EP, Hindes A, Coblitz B, Johnson SL. Mutations in *connexin43* (*GJA1*) perturb bone growth in zebrafish fins. Dev Biol. 2005 Feb 1;278(1):208–19.

37. Hoptak-Solga AD, Klein KA, DeRosa AM, White TW, Iovine MK. Zebrafish short fin mutations in Connexin43 lead to aberrant gap junctional intercellular communication. FEBS Lett. 2007 Jul 10;581(17):3297–302.

38. Harper M, Hu Y, Donahue J, Acosta B, Dievenich Braes F, Nguyen S, et al. Thyroid hormone regulates proximodistal patterning in fin rays. Proc Natl Acad Sci. 2023 May 23;120(21):e2219770120.

39. Cardeira-da-Silva J, Bensimon-Brito A, Tarasco M, Brandão AS, Rosa JT, Borbinha J, et al. Fin ray branching is defined by TRAP+ osteolytic tubules in zebrafish. Proc Natl Acad Sci. 2022 Nov 29;119(48):e2209231119.

40. Wang YT, Tseng TL, Kuo YC, Yu JK, Su YH, Poss KD, et al. Genetic reprogramming of positional memory in a regenerating appendage. Curr Biol CB. 2019 Dec 16;29(24):4193–4207.e4.

41. Litingtung Y, Dahn RD, Li Y, Fallon JF, Chiang C. Shh and Gli3 are dispensable for limb skeleton formation but regulate digit number and identity. Nature. 2002 Aug;418(6901):979–83.

42. Lettice LA, Heaney SJH, Purdie LA, Li L, de Beer P, Oostra BA, et al. A long-range Shh enhancer regulates expression in the developing limb and fin and is associated with preaxial polydactyly. Hum Mol Genet. 2003 Jul 15;12(14):1725–35.

43. Riddle RD, Johnson RL, Laufer E, Tabin C. Sonic hedgehog mediates the polarizing activity of the ZPA. Cell. 1993 Dec 31;75(7):1401–16.

44. Masuya H, Sagai T, Wakana S, Moriwaki K, Shiroishi T. A duplicated zone of polarizing activity in polydactylous mouse mutants. Genes Dev. 1995 Jul 1;9(13):1645–53.

45. Hadzhiev Y, Lele Z, Schindler S, Wilson SW, Ahlberg P, Strähle U, et al. Hedgehog signaling patterns the outgrowth of unpaired skeletal appendages in zebrafish. BMC Dev Biol. 2007 Jun 27;7(1):75.

46. Laforest L, Brown CW, Poleo G, Geraudie J, Tada M, Ekker M, et al. Involvement of the sonic hedgehog, patched 1 and bmp2 genes in patterning of the zebrafish dermal fin rays. Development. 1998 Nov 1;125(21):4175–84.

47. Quint E, Smith A, Avaron F, Laforest L, Miles J, Gaffield W, et al. Bone patterning is altered in the regenerating zebrafish caudal fin after ectopic expression of sonic hedgehog and bmp2b or exposure to cyclopamine. Proc Natl Acad Sci. 2002 Jun 25;99(13):8713–8.

48. Armstrong BE, Henner A, Stewart S, Stankunas K. Shh promotes direct interactions between epidermal cells and osteoblast progenitors to shape regenerated zebrafish bone. Development. 2017 Apr 1;144(7):1165–76.

49. Shen MC, Ozacar AT, Osgood M, Boeras C, Pink J, Thomas J, et al. Heat-shock–mediated conditional regulation of hedgehog/gli signaling in zebrafish. Dev Dyn. 2013;242(5):539–49.

50. Arveseth CD, Happ JT, Hedeen DS, Zhu JF, Capener JL, Shaw DK, et al. Smoothened transduces Hedgehog signals via activity-dependent sequestration of PKA catalytic subunits. PLOS Biol. 2021 Apr 22;19(4):e3001191.

51. AlMuraikhi N, Almasoud N, Binhamdan S, Younis G, Ali D, Manikandan M, et al. Hedgehog Signaling Inhibition by Smoothened Antagonist BMS-833923 Reduces Osteoblast Differentiation and Ectopic Bone Formation of Human Skeletal (Mesenchymal) Stem Cells. Stem Cells Int. 2019 Nov 21;2019:3435901.

52. Lin TL, Matsui W. Hedgehog pathway as a drug target: Smoothened inhibitors in development. OncoTargets Ther. 2012 Dec 31;5:47–58.

53. Salis P, Roux N, Huang D, Marcionetti A, Mouginot P, Reynaud M, et al. Thyroid hormones regulate the formation and environmental plasticity of white bars in clownfishes. Proc Natl Acad Sci U S A. 2021 Jun 8;118(23):e2101634118.

54. Randall JE, Kuiter RH. Three New Labrid Fishes of the Genus Coris from the Western Pacific. Pac Sci. 1982 Apr;36(2).

55. Fricke R, Durville P. Coris flava, a new deep water species of wrasse from La Réunion, southwestern Indian Ocean (Teleostei: Labridae). FishTaxa. 2021 Dec 24;22:23–36.

56. Tanaka Y, Okayama S, Ansai S, Abe G, Tamura K. Fin elaboration via anterior-posterior regulation by Hedgehog signaling in teleosts. bioRxiv; 2023. p. 2023.10.10.557878.

57. Placinta M, Shen MC, Achermann M, Karlstrom RO. A laser pointer driven microheater for precise local heating and conditional gene regulation in vivo. Microheater driven gene regulation in zebrafish. BMC Dev Biol. 2009 Dec 30;9:73.

58. Bouldin CM, Kimelman D. Dual Fucci: A New Transgenic Line for Studying the Cell Cycle from Embryos to Adults. Zebrafish. 2014 Apr 1;11(2):182–3.

59. Goldsmith MI, Iovine MK, O’Reilly-Pol T, Johnson SL. A developmental transition in growth control during zebrafish caudal fin development. Dev Biol. 2006 Aug 15;296(2):450–7.

60. Huang P, Xiong F, Megason SG, Schier AF. Attenuation of Notch and Hedgehog Signaling Is Required for Fate Specification in the Spinal Cord. PLoS Genet. 2012 Jun 7;8(6):e1002762.

61. Silic MR, Wu Q, Kim BH, Golling G, Chen KH, Freitas R, et al. Potassium Channel-Associated Bioelectricity of the Dermomyotome Determines Fin Patterning in Zebrafish. Genetics. 2020 Aug 1;215(4):1067–84.

62. Chiang C, Litingtung Y, Lee E, Young KE, Corden JL, Westphal H, et al. Cyclopia and defective axial patterning in mice lacking Sonic hedgehog gene function. Nature. 1996 Oct;383(6599):407–13.

63. Zhulyn O, Nieuwenhuis E, Liu YC, Angers S, Hui C chung. Ptch2 shares overlapping functions with Ptch1 in Smo regulation and limb development. Dev Biol. 2015 Jan 15;397(2):191–202.

64. Sagai T, Masuya H, Tamura M, Shimizu K, Yada Y, Wakana S, et al. Phylogenetic conservation of a limb-specific, cis-acting regulator of Sonic hedgehog (Shh). Mamm Genome. 2004 Jan 1;15(1):23–34.

65. Yang Y, Drossopoulou G, Chuang PT, Duprez D, Marti E, Bumcrot D, et al. Relationship between dose, distance and time in Sonic Hedgehog-mediated regulation of anteroposterior polarity in the chick limb. Development. 1997 Nov 1;124(21):4393–404.

66. Harfe BD, Scherz PJ, Nissim S, Tian H, McMahon AP, Tabin CJ. Evidence for an Expansion-Based Temporal Shh Gradient in Specifying Vertebrate Digit Identities. Cell. 2004 Aug 20;118(4):517–28.

67. Dahn RD, Davis MC, Pappano WN, Shubin NH. Sonic hedgehog function in chondrichthyan fins and the evolution of appendage patterning. Nature. 2007 Jan;445(7125):311–4.

68. Onimaru K, Kuraku S, Takagi W, Hyodo S, Sharpe J, Tanaka M. A shift in anterior–posterior positional information underlies the fin-to-limb evolution. Bronner ME, editor. eLife. 2015 Aug 18;4:e07048.

69. Letelier J, de la Calle-Mustienes E, Pieretti J, Naranjo S, Maeso I, Nakamura T, et al. A conserved Shh cis-regulatory module highlights a common developmental origin of unpaired and paired fins. Nat Genet. 2018 Apr;50(4):504–9.

70. Letelier J, Naranjo S, Sospedra-Arrufat I, Martinez-Morales JR, Lopez-Rios J, Shubin N, et al. The *Shh* / *Gli3* gene regulatory network precedes the origin of paired fins and reveals the deep homology between distal fins and digits. Proc Natl Acad Sci. 2021 Nov 16;118(46):e2100575118.

71. Hawkins MB, Jandzik D, Tulenko FJ, Cass AN, Nakamura T, Shubin NH, et al. An Fgf–Shh positive feedback loop drives growth in developing unpaired fins. Proc Natl Acad Sci. 2022 Mar 8;119(10):e2120150119.

72. Neumann CJ, Grandel H, Gaffield W, Schulte-Merker S, Nüsslein-Volhard C. Transient establishment of anteroposterior polarity in the zebrafish pectoral fin bud in the absence of sonic hedgehog activity. Development. 1999 Nov 1;126(21):4817–26.

73. Chiang C, Litingtung Y, Harris MP, Simandl BK, Li Y, Beachy PA, et al. Manifestation of the Limb Prepattern: Limb Development in the Absence of Sonic Hedgehog Function. Dev Biol. 2001 Aug 15;236(2):421–35.

74. Zákány J, Kmita M, Duboule D. A Dual Role for Hox Genes in Limb Anterior-Posterior Asymmetry. Science. 2004 Jun 11;304(5677):1669–72.

75. Lewandowski JP, Du F, Zhang S, Powell MB, Falkenstein KN, Ji H, et al. Spatiotemporal regulation of GLI target genes in the mammalian limb bud. Dev Biol. 2015 Oct 1;406(1):92– 103.

76. Nakamura T, Gehrke AR, Lemberg J, Szymaszek J, Shubin NH. Digits and fin rays share common developmental histories. Nature. 2016 Sep;537(7619):225–8.

77. Towers M, Mahood R, Yin Y, Tickle C. Integration of growth and specification in chick wing digit-patterning. Nature. 2008 Apr;452(7189):882–6.

78. Zhu J, Nakamura E, Nguyen MT, Bao X, Akiyama H, Mackem S. Uncoupling Sonic Hedgehog Control of Pattern and Expansion of the Developing Limb Bud. Dev Cell. 2008 Apr 15;14(4):624–32.

79. Lopez-Rios J, Speziale D, Robay D, Scotti M, Osterwalder M, Nusspaumer G, et al. GLI3 Constrains Digit Number by Controlling Both Progenitor Proliferation and BMP-Dependent Exit to Chondrogenesis. Dev Cell. 2012 Apr 17;22(4):837–48.

80. Boehm B, Westerberg H, Lesnicar-Pucko G, Raja S, Rautschka M, Cotterell J, et al. The Role of Spatially Controlled Cell Proliferation in Limb Bud Morphogenesis. PLoS Biol. 2010 Jul 13;8(7):e1000420.

81. Lauder GV. Caudal Fin Locomotion in Ray-finned Fishes: Historical and Functional Analyses1. Am Zool. 1989 Feb 1;29(1):85–102.

82. Lauder GV. Function of the Caudal Fin During Locomotion in Fishes: Kinematics, Flow Visualization, and Evolutionary Patterns1. Am Zool. 2000 Feb 1;40(1):101–22.

83. Davesne D, Friedman M, Schmitt AD, Fernandez V, Carnevale G, Ahlberg PE, et al. Fossilized cell structures identify an ancient origin for the teleost whole-genome duplication. Proc Natl Acad Sci. 2021 Jul 27;118(30):e2101780118.

84. Song J, Zhong Y, Du R, Yin L, Ding Y. Tail shapes lead to different propulsive mechanisms in the body/caudal fin undulation of fish. Proc Inst Mech Eng Part C J Mech Eng Sci. 2021 Jan 1;235(2):351–64.

85. Arratia G. Complexities of Early Teleostei and the Evolution of Particular Morphological Structures through Time. Copeia. 2015 Dec;103(4):999–1025.

86. Thieme P, Warth P, Moritz T. Development of the caudal-fin skeleton reveals multiple convergent fusions within Atherinomorpha. Front Zool. 2021 Apr 26;18(1):20.

87. Hoshino K. Homologies of the caudal fin rays of Pleuronectiformes (Teleostei). Ichthyol Res. 2001 Aug 1;48(3):231–46.

88. DeLaurier A, Eames BF, Blanco-Sánchez B, Peng G, He X, Swartz ME, et al. Zebrafish sp7:EGFP: a transgenic for studying otic vesicle formation, skeletogenesis, and bone regeneration. Genes N Y N 2000. 2010 Aug;48(8):505–11.

89. Kirby BB, Takada N, Latimer AJ, Shin J, Carney TJ, Kelsh RN, et al. In vivo time-lapse imaging shows dynamic oligodendrocyte progenitor behavior during zebrafish development. Nat Neurosci. 2006 Dec;9(12):1506–11.

90. Bouldin CM, Snelson CD, Farr GH, Kimelman D. Restricted expression of cdc25a in the tailbud is essential for formation of the zebrafish posterior body. Genes Dev. 2014 Feb 15;28(4):384–95.

91. van Eeden FJM, Granato M, Schach U, Brand M, Furutani-Seiki M, Haffter P, et al. Genetic analysis of fin formation in the zebrafish, Danio rerio. Development. 1996 Dec 1;123(1):255–62.

92. Walker M, Kimmel C. A two-color acid-free cartilage and bone stain for zebrafish larvae. Biotech Histochem. 2007 Jan 1;82(1):23–8.

93. Schindelin J, Arganda-Carreras I, Frise E, Kaynig V, Longair M, Pietzsch T, et al. Fiji: an open-source platform for biological-image analysis. Nat Methods. 2012 Jul;9(7):676–82.

